# Predicting Global Forest Reforestation Potential

**DOI:** 10.1101/210062

**Authors:** Thomas W. Crowther, Henry Glick, Daniel Maynard, Will Ashley-Cantello, Tom Evans, Devin Routh

## Abstract

Forests are important determinants of the carbon cycle, and they provide countless ecosystem services to support billions of people worldwide. Global-scale forest restoration is one of our most effective weapons in the fight against biodiversity loss, rural poverty and climate change. In this report, we generate a spatial map of tree density within the potential forest restoration areas delineated by the IUCN/WRI’s “Atlas of Forest Landscape Restoration Opportunities” to estimate the potential number of trees that could be restored at a global scale. We also estimate the number of trees that might be saved by avoiding deforestation in currently forested areas.

We show that the restoration areas have the capacity to support a total of 1.33 trillion trees. However, given that a considerable proportion of these areas already contain forests, we estimate that 589 billion new trees (larger than 10 cm diameter) could be restored within these areas, which would have the potential to store 65–91 Gigatonnes of carbon after reaching forest maturation. These values will increase marginally over time, as deforestation is responsible for the removal of living trees within the restoration areas. However, if only 50% or 25% of the mosaic areas (the largest of the designated restoration types) are available for reforestation, this total number will fall to approximately 360, or 246 billion trees, respectively, with corresponding decreases in potential carbon storage. Given that anthropogenic carbon emissions are currently in the order of 9–12 Gigatonnes per year, effective global-scale restoration might potentially have a valuable impact on global-scale climate mitigation over the rest of this century.

This report was produced with funding from WWF-UK as part of the Trillion Trees programme with the Wildlife Conservation Society and BirdLife International

## I. Introduction

Given the vast array of ecosystem services provided by trees, the restoration of forests at a global scale represents a valuable approach for improving biodiversity, air quality and human health[1][2] In particular, the unique capacity of trees to absorb *CO*_2_ directly from the atmosphere makes forest restoration a viable method to partially mitigate the effects of anthropogenic climate change. However, generating meaningful restoration targets requires an understanding of the current extent of forest trees and areas available for potential restoration.

A growing body of research is enabling us to generate an understanding of forests at a global scale. This global forest system covers 3.9 billion hectares. It contains approximately 3.04 trillion trees (over 10 cm DBH),[2] which store approximately 250–350 Gt of carbon. [4][5][6] However, little is known about the potential for forest restoration in currently un-forested regions. Until now, it has remained unclear how many trees can potentially be restored on Earth, and how much carbon might they store. Here, we use the predictive equations that were generated in the first analysis of global tree density,[2] to predict potential tree numbers and their distributions within the ‘restoration areas’ as defined by the IUCN/WRI’s ‘Atlas of Forest Landscape Restoration Opportunities’.[7] Using existing estimates of forest biomass,[4][5][6] we then approximate how much carbon these trees could potentially store if they were restored to equivalent extents as seen in current existing forests. We also use projections of future deforestation (provided by the WWF Living Forests Report [8]) to estimate how potential restoration numbers might be affected by deforestation under different future land use scenarios. This understanding of global deforestation rates can help to place this restoration information into context, and allow us to understand how forest restoration might influence net changes in tree numbers and carbon storage on a global scale.

## II. RESULTS & DISCUSSION

### i. Projected global deforestation rates

Before calculating potential restoration estimates, it is important for us to consider the current and on-going rates of deforestation that will place these results into context. The current net global rates of forest loss have been estimated at approximately 10 billion trees each year.[2] The WWF’s ‘Living forests report’[8] allows us to project these total losses into the future by providing estimates of global forest area loss under various different global deforestation scenarios over the next 35 years. Based on these deforestation estimates, the starting rates of tree loss expected in 2005 (10 billion trees per year) are approximately equivalent the current deforestation rate. If the global rates of forest loss remained consistent, we would expect to lose approximately 350 billion trees by 2050. However, deforestation rates are then expected to fall drastically over the next 3 decades. Even under the business as usual (do nothing) scenario, rates of forest loss are expected to gradually decline, so that total cumulative losses are approximately 118 billion by 2050 (see Figure 1 for projected deforestation rates under each of the potential scenarios). More aggressive reductions in global deforestation rates in the short term can potentially allow us to avoid the majority of these tree losses, with only 5.9 billion trees estimated to be lost under the most optimistic (unlikely) ‘Target met’ scenario (Table 1).

**Figure 1:**
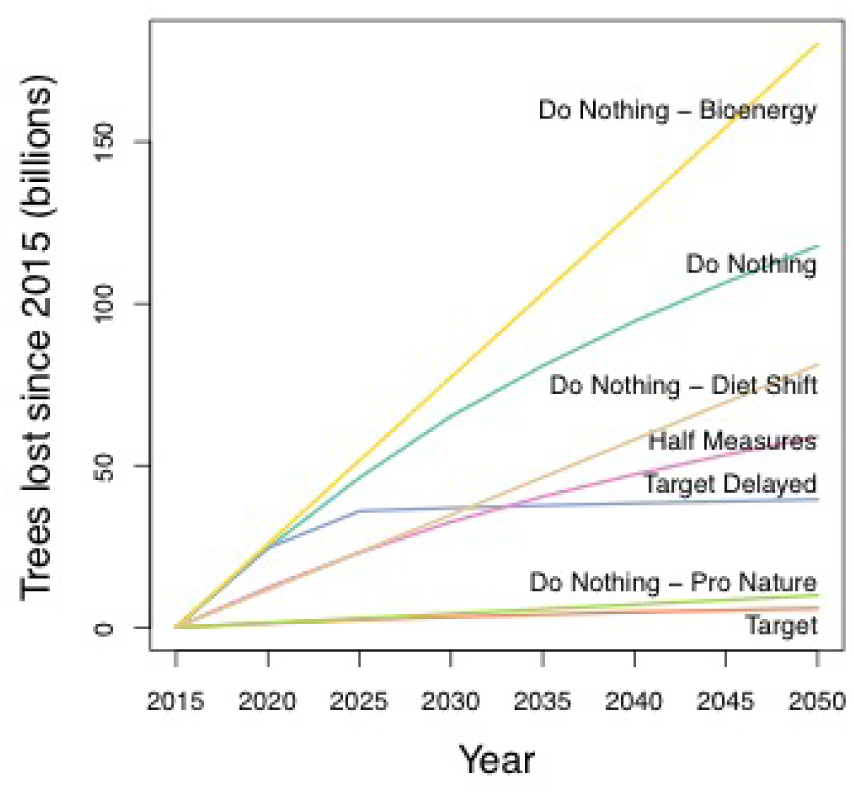
*Global projections of cumulative trees lost by 2050 under different deforestation scenarios projected by the WWF’s living trees report.[3]*

**Table 1:** *Summary of the total potential trees expected to be lost over the next 35 years, and the total number of trees that could be restored globally at present.*

| Billions of trees lost/added |  |  |  |  |  |  |  |
| --- | --- | --- | --- | --- | --- | --- | --- |
| Deforestation scenarios | Do Nothing | Target met (ZNDD 2020) | Target delayed (ZNDD 2030) | Half measures | Pro nature | Bioenergy | Diet Shift |
| 2040 | 65.2 | 1.2 | 36.9 | 32.6 | 4.2 | 77.2 | 34.8 |
| 2030 | 94.5 | 4.7 | 38.3 | 47.3 | 7.1 | 128.7 | 58.0 |
| 2050 | 117.7 | 5.9 | 39.5 | 58.9 | 9.9 | 180.2 | 81.1 |
| Restoration scenarios | High: Maximise tree cover in all areas |  | Medium: 'Widescale' and 'Remote' areas at 100%, 'Mosaics' at 50% tree cover |  | Low: 'Widescale' and 'Remote' areas at 100%, 'Mosaics' at 25% tree cover. |  |  |
| 40cm DBH | 7661.1 |  | 4691.8 |  | 4691.8 |  |  |
| 10cm DBH | 589.2 |  | 360.8 |  | 360.8 |  |  |
| 1cm DBH | 125.7 |  | 77.4 |  | 77.4 |  |  |

**Table 2:** *Mean current tree densities for each of the world’s forested biomes.*

| Biome | Mean tree density(trees per hectare) |
| --- | --- |
| Boreal Forests/Taiga | 967.59 |
| Deserts and Xeric Shrublands | 244.57 |
| Flooded Grasslands and Savannas | 497.24 |
| Mangroves | 494.08 |
| Mediterranean Forests, Woodlands and Scrub | 881.39 |
| Montane Grasslands and Shrublands | 799.31 |
| Temperate Broadleaf and Mixed Forests | 486.41 |
| Temperate Conifer Forests | 426.19 |
| Temperate Grasslands, Savannas and Shrublands | 285.47 |
| Tropical and Subtropical Coniferous Forests | 426.19 |
| Tropical and Subtropical Dry Broadleaf Forests | 372.35 |
| Tropical and Subtropical Grasslands, Savannas and Shrublands | 292.88 |
| Tropical and Subtropical Moist Broadleaf Forests | 779.26 |
| Tundra | 1042.03 |

At the biome-level, the areas that are most susceptible to wide-scale tree loss are the ‘tropical and subtropical moist broadleaf forests’ and the ‘boreal forests’, which are expected to experience 37 and 35% of the global tree loss, respectively. With considerable areas of undamaged, old-growth forests these regions are among the most valuable from biodiversity and a carbon storage perspectives, but they also provide the most attractive opportunities for logging and land conversion. Given the nature of global land use projections into the future, the deforestation projections are not spatially explicit. As such, we are unable to estimate potential tree losses at a higher spatial resolution.

The deforestation scenarios that we present above are based on the WWF’s Living Forests Report, which projected considerable reductions in deforestation rates between the years of 2005 and 2015. These optimistic expectations have not, however, been realized. Indeed, the predicted loss rate for 2005 is approximately equivalent to present day losses, and projected loss rates for 2015 are considerably lower. As such, the expected losses (above) are considerably lower than the realistic ones. To account for this, we re-ran the analysis, but with the starting deforestation rates being equal to present day rates. This effectively involved shifting the starting date forward by 10 years, so that the 2005 rates are equivalent to the 2015 rates. This analysis revealed considerably greater tree losses under all scenarios. In particular, the final losses by 2050 for the ‘do nothing âĂŞ bioenergy’, ‘do nothing’, ‘target delayed’ and ‘target’ scenarios increase to 179 billion, 157 billion, 94 billion and 54 billion trees, respectively (See figure 2 for updated estimates of deforestation).

**Figure 2:**
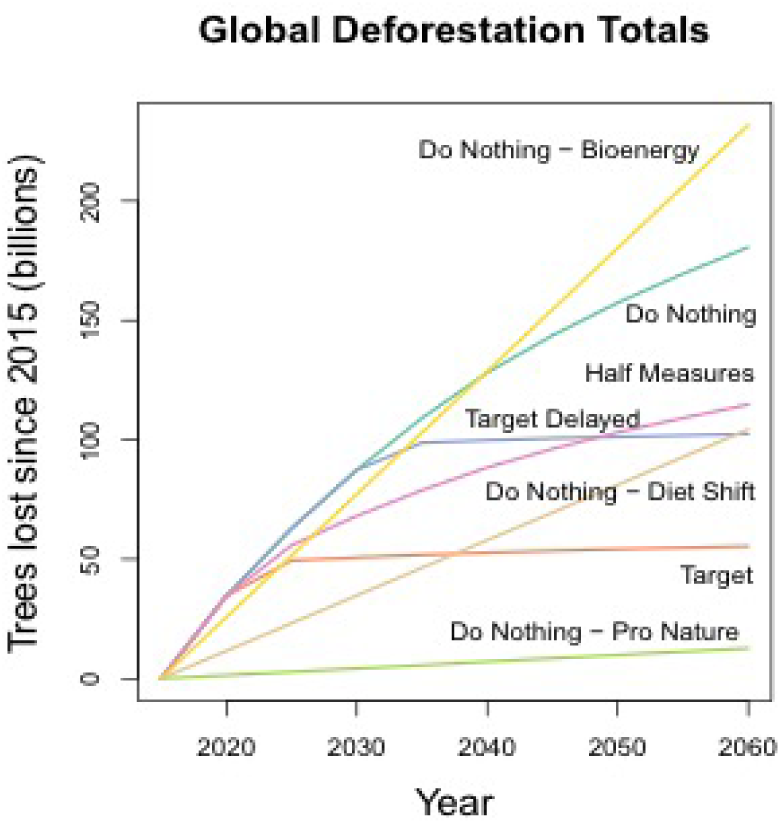
*Altered global projections of cumulative trees lost by 2050 under different deforestation scenarios projected by the WWF’s living trees report.[3] These estimates force the starting rates of deforestation to match the currently observed rates.*

### ii. Total potential tree numbers that can fit into the restoration areas

The Atlas of Forest Landscape Restoration Opportunities [7] delineates three specific categories of potential restoration area: ‘widescale’, ‘mosaic’ and ‘remote’. Based on their descriptions, ‘widescale’ refers to regions that could be completely reforested, ‘mosaic’ refers to regions that could be partially reforested, and ‘remote’ describes the areas that are available for complete restoration but they exist within remote and challenging areas.

Adding together the tree count currently existing in living forests within the restoration areas and the potential for new trees (using the currrent densities in those areas), we estimate that approximately 1.33 trillion trees (≥ 10 cm trunk diameter at breast height) could exist within the global restoration areas. Of these trees, 309 billion, 867 billion and 155 billion trees could exist within wide-scale, mosaic and remote areas, respectively. However, this does *not* represent the total number of trees that can be restored within these areas because large areas within these restoration maps already contain healthy forests.

We therefore estimated the number of trees that could be restored within the currently non-forested areas under each restoration scenario outlined in IUCN/WRI’s ‘Atlas of Forest Landscape Restoration Opportunities’. [7] Excluding the trees that currently exist, our model reveals that an additional 589 (±55) billion trees (≥10 cm trunk diameter) could be restored at present global tree densities if 100% of the available restoration land is reforested (see Figure 3 for a breakdown of the potential number of trees to be restored within each of 14 terrestrial biomes that contain forests). Of these, approximately 99 billion, 457 billion and 34 billion would exist within widescale, mosaic and remote restoration areas, respectively.

**Figure 3:**
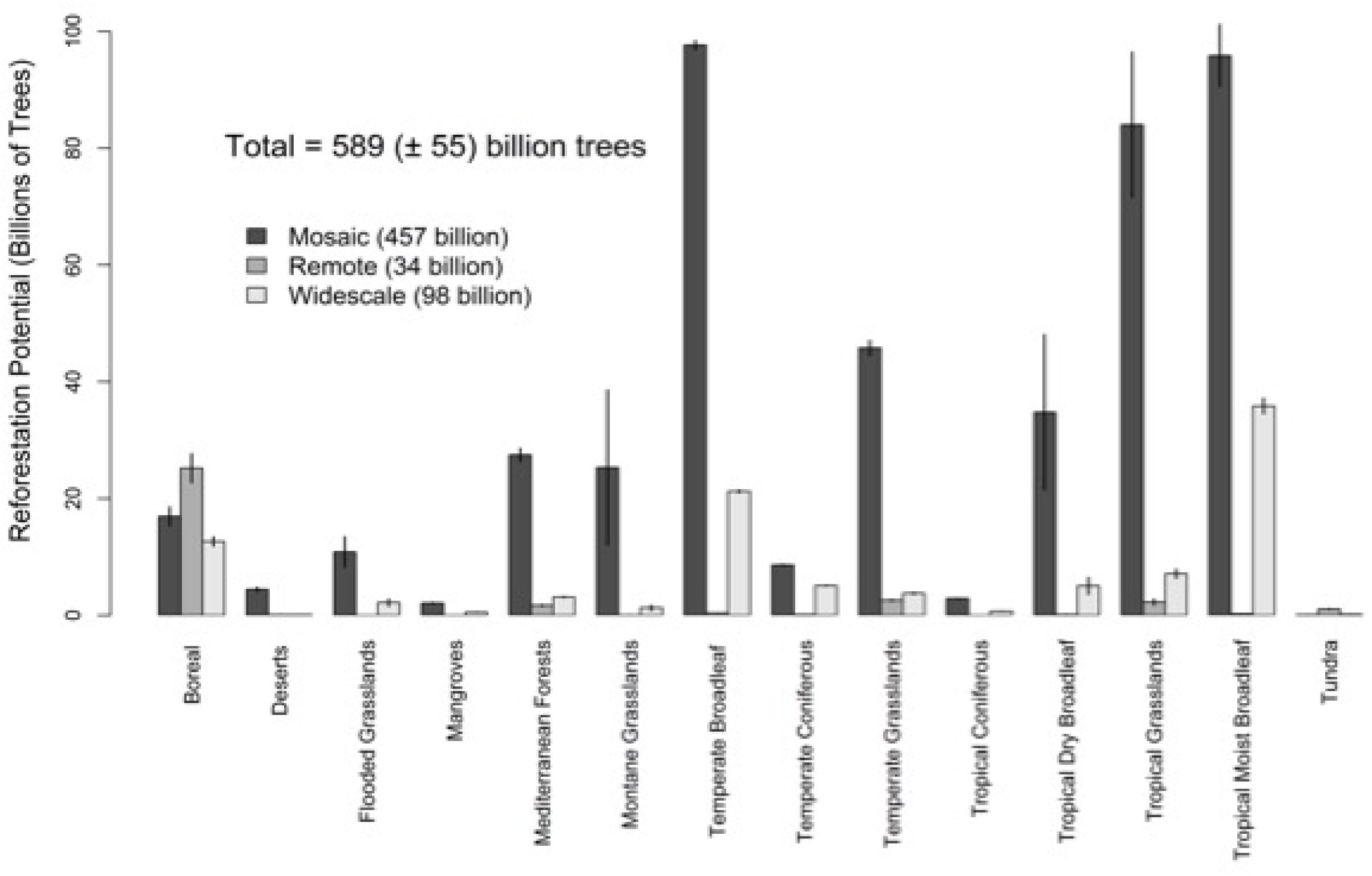
*Breakdown of reforestation potential within the global restoration areas. Shows total numbers (and the 95% confidence intervals) of trees (≥10 cm diameter) that could be restored within ‘mosaic’, ‘remote’, and ‘widescale’ restoration areas, within each of the 14 forested biomes (delineated by the Nature Conservancy). To estimate the total numbers if only 50 or 25% of the areas are available for deforestation, just divide any of these numbers by 2 or 4.*

These numbers assume complete restoration of targeted areas, whereas the total number of actual trees will depend on the proportion of land that is realistically available for reforestation. The Atlas of Forest Landscape Restoration Opportunities [7] delineates the areas where human activity are most likely to encroach on potential forest land, and they are contained within the ‘Mosaic’ restoration areas. If only 50% or 25% of the mosaic areas are available for reforestation, the total number of trees for restoration falls to approximately 360, or 246 billion, respectively (see Table 1). In addition, a number of the sites highlighted for potential restoration include grassland sites that would not naturally support forests. We would highlight that conserving local biodiversity will often require that reforestation does not occur in those areas.

Ongoing deforestation is expected to reduce the number of trees that currently exist within these restoration areas. As a result, the area available for potential reforestation is likely to increase over time. Based on projected deforestation rates under the different land-use scenarios,[8] we estimate that the total number of trees to be restored within the global restoration zones is likely to increase marginally over the next 35 years (see Figure 4 for estimates of how the total reforestation numbers will increase under the different land-use scenarios).

**Figure 4:**
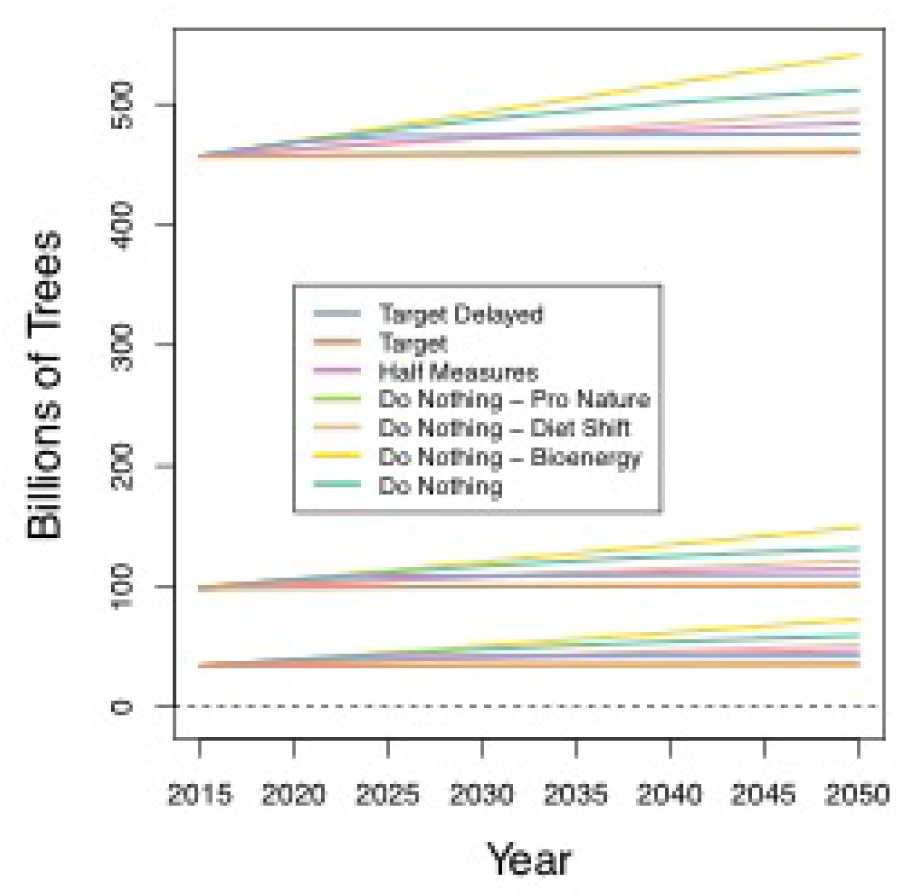
*Impacts of deforestation within the IUCN/WRI restoration areas on total reforestation potential (total trees 10 cm diameter) within the UICN/WRI restoration areas. Under several land use scenarios, [8] the current areas of forest within the restoration areas will continue to decline over the next 35 years. The area of available reforestation land (and the total potential reforestation) will, therefore, increase marginally over this period.*

**Figure 5:**
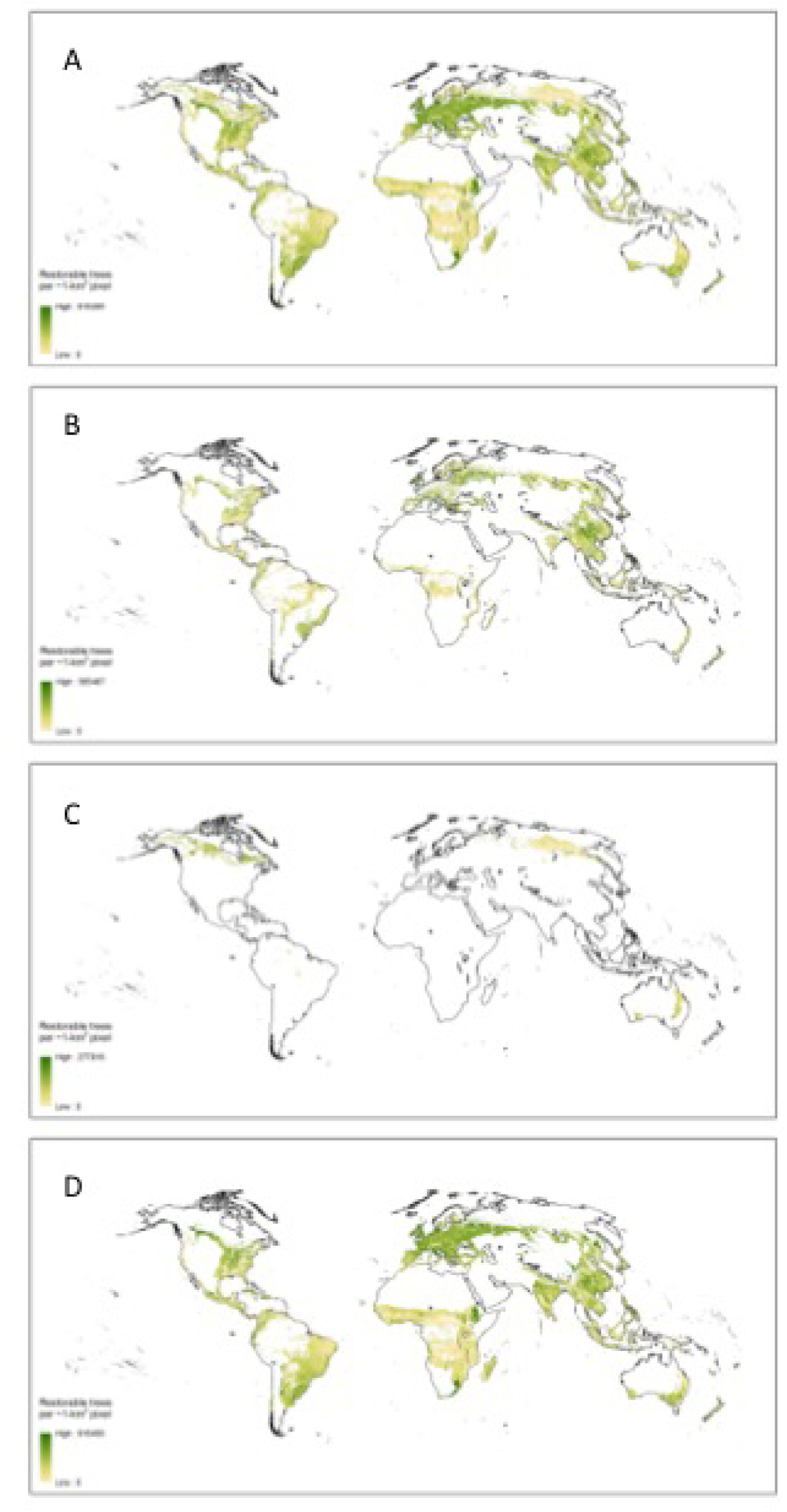
Maps of global forest restoration potential in terms of tree density. Panel ‘A’ shows the total restoration potential across all restoration types. Panels ‘B’, ‘C’, and ‘D’ show the potential restoration in Widescale, Remote, and Mosaic restoration regions as delineated by the IUCN/WRI’s ‘Atlas of Forest Landscape Restoration Opportunities’.[7].

### iii. Quantifying the global impact

If trees were restored across 100% of the global restoration areas, the impact would represent an increase of approximately 19.36% on top of the current global number. If 50% of the mosaic restoration areas were restored, this percentage would still represent a considerable proportion (approximately 11.87%) of the global tree number.

If we assume that these trees were able to store an equivalent amount of carbon that currently exists within equivalent forested regions,[4][5][6] then it is possible to estimate how much carbon might potentially be stored in these restoration regions (See Table 3). Based on this information we estimate that, with 100% restoration across the entire restoration area, it might be possible to for the trees to store 65–91 Gigatonnes of carbon in their aboveground and belowground biomass (see Table 3). In contrast, if only 50% of the Mosaic restoration areas are restored, these potential carbon storage falls to 36–50 Gigatonnes. The average age of the worlds extant forests is unclear so it is difficult to determine how long it might take for these carbon stocks to accumulate. But based on our understanding of global forest systems, we estimate that it would be likely to take at least 50–100 for carbon stocks of this magnitude to be restored within forest ecosystems

**Table 3:**
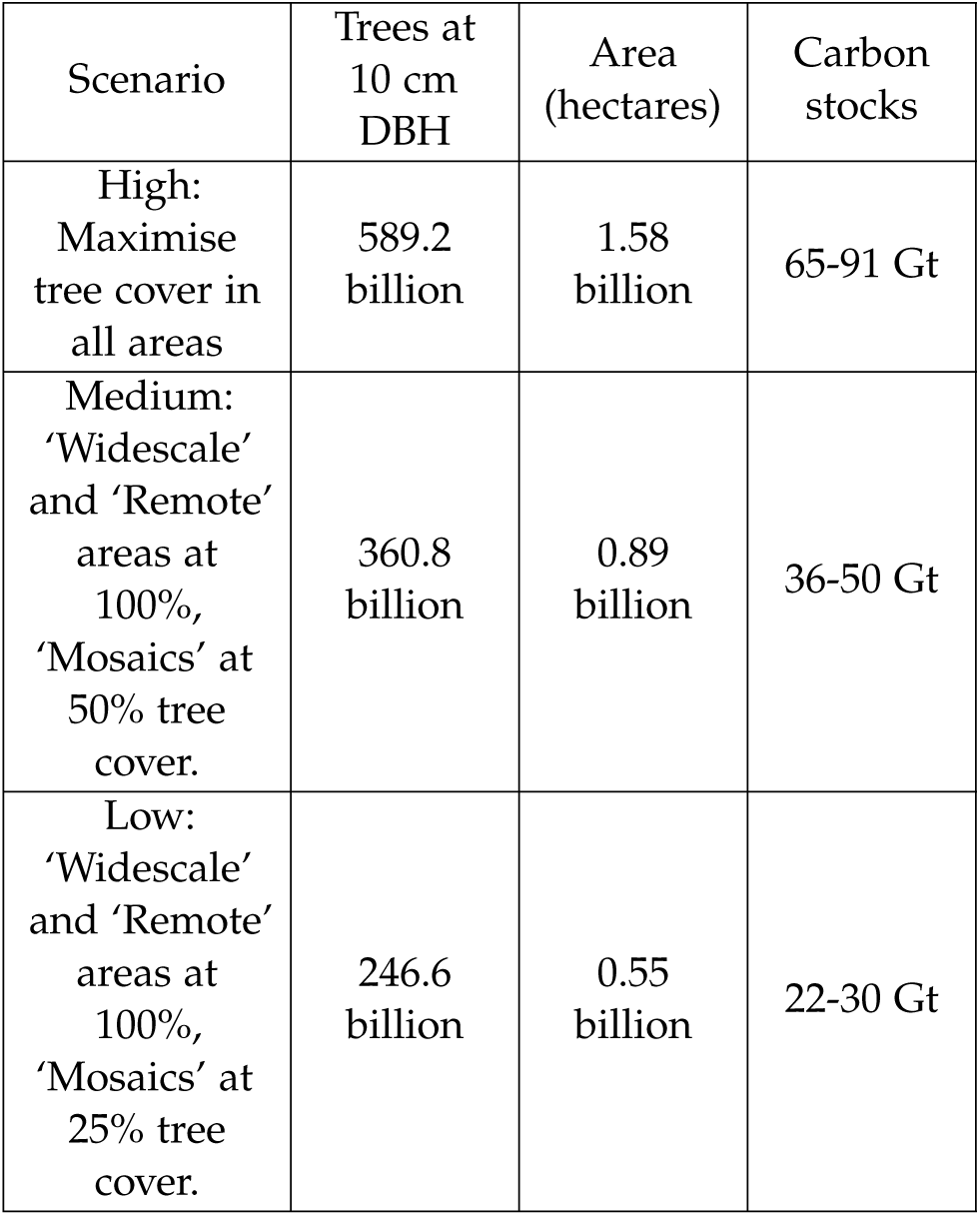
*Summary of the restoration area, and potential carbon storage for each of the restoration scenarios outlined in the IUCN/WRI’s ‘Atlas of Forest Landscape Restoration Opportunities’[7]. See supplementary Table S1 for a full breakdown of these values by country.*

### iv. Tree density changes over time

Following almost all of the National Forest Inventories around the world, most of our analyses focus on plants with woody stems larger than 10 cm diameter at breast height. However, in the short-term, the trees restored or regenerating naturally in any global reforestation efforts will be considerably smaller than this. Relationships between tree density and stem diameter reveal that the densities of smaller trees are always considerably higher than those of larger trees. Based on these relationships, we estimate that at least 7.66 trillion saplings (trees with 1 cm diameter) could be restored or regenerated within the non-forested portions of the restoration areas. However, if only 50% or 25% of the Mosaic areas are available for restoration, this number falls to 4.7 and 3.0 trillion, respectively (see Table 1). If these restored forests remained undisturbed until maturity, the resulting number of mature trees (individuals larger than 40 cm diameter) would be approximately 125 billion, 77 billion and 50 billion for 100% 50% and 25% remote restoration scenarios, respectively.

### v. Limitations and considerations

The >420,000 plot measurements that informed our spatial models represent current tree densities. These plot locations include a vast number of measurements from forests that have been heavily impacted by humans. Across every terrestrial biome (except montane grasslands), human activity has consistently reduced tree density. Therefore, the tree densities that we project here are likely to be lower than the potential tree densities. For example, the mean tree density that we report within boreal forest areas is approximately 968 trees/ha. This equates to 9.68 trees for each 10*m*^2^ area of forest. Given that tree density measurements collected in the global tree density study reached 10 times this value, it is likely that potential tree numbers could exceed those that we present here.

Despite the strength of our model predictions at large spatial scales, the accuracy of our estimates is constrained by the robustness of the estimates in the WWF’s Living Forest Report, and the Atlas of Forest Landscape Restoration Opportunities.[7] Firstly, the projected rates of deforestation in the Living Forests Report are highly optimistic, with considerable reductions expected over time (even under the business as usual scenario). This temporal extrapolation can be lead to uncertainty in our projections. If these expectations are not realized then the real global tree losses are likely to be considerably higher than we present.

Secondly, although the Atlas of Forest Landscape Restoration Opportunities [7] is considerably more detailed and spatially explicit, the global-scale of these estimates also limits confidence in these estimates. Given the size of these restoration areas and the scale of the map, it is likely that restoration areas exist in smaller areas outside of the predicted zones. Similarly, considerable proportions of the restoration areas will not be suitable for reforestation. The restoration zones contain urban and agricultural land as well as considerable areas of natural grassland that should be maintained if we intend to conserve local biodiversity. In addition we have shown that a considerable proportion of these restoration areas already contain forests and are not likely to be suitable for future reforestation. However, these estimates serve as an illustrative set of boundaries for the potential for future global restoration potential.

### vi. Potential future work

Given the importance of the WWF’s Living Forests Report,[8] and the Atlas of Forest Landscape Restoration Opportunities [7] for projecting potential changes in tree numbers, future work should focus on evaluating and refining these estimates. The Atlas of Forest Landscape Restoration Opportunities [7] represents the most recent global resource to guide future restoration decisions. However, it is thematically coarse. By simply highlighting areas that might be eligible for future restoration, the report is unable to differentiate between different land use types. Clearly there are considerable portions of land within the designated restoration types that will not be available for reforestation. For example, many natural grasslands and deserts should not be considered suitable for reforestation. Developing more fine-scale differentiation between different land uses within the restoration areas should be a research priority over the next 5 years. Satellite-based remote sensing can serve as a valuable resource for differentiating between desert, grassland, existing forest and urban land. Following this, it is necessary to identify the publicly available land that is not currently being used for agriculture. Spatially explicit maps of private land can serve as a guideline for identifying available land. Once the areas for potential reforestation are more tightly delineated, the remaining maps can be paired with information about tree densities to drastically improve the confidence of potential reforestation projections.

The WWF’s projections of future forest loss have already not been met, highlighting that the future projected estimates are likely to be unrealistic. Evaluating these estimates in light of recent changes in global deforestation rates (as highlighted by Hansen *et al.*[3]) should be an essential short-term (1 year) challenge to improve the effectiveness of future restoration efforts.

## III. MATERIALS & METHODS

The methods detailed here for estimation of forest tree density are equivalent to many of those first outlined in Crowther et al,[2] and a detailed explanation can be found in Glick *et al.*[9]. Portions of the following text have been adapted from the original work and the reader is referred to the original publication for additional detail.

### i. Data collection and standardization

Plot-level data had already been collected from international forestry databases, including the Global Index of Vegetation-Plot Database (GIVD http://www.givd.info), the Smithsonian Tropical Research Institute (http://www.stri.si.edu), ICP-Level-I plot data which covers most of Europe (http://www.icpforests.org), and National Forest Inventory (NFI) analyses from 21 countries, including the USA (http://fia.fs.fed.us/) and Canada (https://nfi.nfis.org/index.php). This information was supplemented with data from peerreviewed studies reporting large international inventories published in the last 10 years (collected using ISI Web of Knowledge, Google Scholar and secondary references).

We included density estimates where individual trees met the criterion of ≥10 cm diameter at breast height (DBH). Although NFI databases can vary slightly in their definition of a mature tree (for example, the US Forest Service Forest Inventory and Analysis (FIA) defines a tree as a plant with woody stems larger than 12.7 cm DBH) the vast majority of sources use 10 cm as the DBH cut-off. Indeed, this was the only size class provided by all broad-scale inventories (including the FIA), so density estimates at other DBH values were excluded. This provided a total of 429,775 measurements of forest tree density (each generated at the hectare scale) that were then linked to spatially explicit remote-sensing data and Geographic Information System (GIS) variables to explore patterns in forest tree density at a global scale.

### ii. Acquisition and preprocessing of spatial data

For predictive model development, we selected 20 geospatial covariates from a larger pool of potential covariates based on uniqueness, spatial resolution and ecological relevance. Covariates were derived through satellite-based remote sensing and ground-based weather stations, and can be loosely grouped into one of four categories: topographic, climatic, vegetative or anthropogenic. Topographic covariates included elevation, slope, aspect (as northness and eastness), latitude (as absolute value of latitude) and a terrain roughness index (TRI). Climatic covariates included annual mean temperature, temperature annual range, annual precipitation, precipitation of driest month, precipitation seasonality (coefficient of variation), precipitation of driest quarter, potential evapotranspiration per hectare per year, and indexed annual aridity. Vegetative covariates included enhanced vegetation index (EVI), leaf area index (LAI), dissimilarity of EVI, contrast of EVI, and angular second moment of EVI (see http://earthenv.org for details). We also included a single anthropogenic covariate: proportion of urban and/or developed land cover.

To account for broad-scale differences in vegetation types, we developed spatial models at the biome scale. Individual predictive models were generated within each of 14 broad ecosystem types (as delineated by the Nature Conservancy’s map of Terrestrial Ecoregions, http://www.nature.org) to improve the accuracy of estimates.

### iii. Statistical modeling

We used generalized linear models with a negative binomial error structure to generate predictive maps of tree numbers within forested ecosystems for each biome. These models were originally generated to predict current worldwide tree densities.[2] Due to the inherently interactive nature of climate, topographic and human impact factors across the globe, we predicted that there would be pronounced non-independence within the full suite of biophysical variables extracted from the compiled GIS layers. To account for this collinearity, we performed ascendant hierarchical clustering using the *hclustvar* function in R’s ClustOfVar package for each biome-level model. This analysis splits the variables into different clusters (similar to principal components) in which all variables correlate with one another. A single best ‘indicator’ variable is then selected from each cluster, based on squared loading values representing the correlation with the central synthetic variable of each cluster (that is, the first principal component of a PCAmix analysis). This set of ‘best’ indicator variables for each biome was then included in all subsequent models used to estimate controls on forest tree density.

Using the resulting set of variables, we derived covariates, coefficients, and variance-covariance matrices for biome-level models through weighted model averaging (see [2] for details). Our modelling approach was used to generate spatially-explicit predictive estimates of potential forest tree density for each square-kilometer pixel within each restoration area. These estimates were subsequently scaled based on the proportion of area within each square-kilometer pixel that is currently un-forested [2] (see spatial modeling for details).

### iv. Spatial modeling

We applied the final negative binomial regression equations to pixel-level spatial data for each potential reforestation zone. For covariates whose values were highly dependent on current numbers of trees or other vegetative characteristics (e.g., vegetative indices), we calculated the mean covariate values among those plots within each biome that were *≥*75% forested. These mean covariate values were then assigned to each potential restoration pixel in each respective biome. This approach allowed us to artificially simulate the vegetative characteristics of forest in areas where there is not currently forest, thus allowing us to model the maximal number of trees that could exist within each fully forested pixel.

Regressions were run in a map algebra framework wherein equation intercepts and coefficients were applied independently to each pixel of our coregistered global covariates to produce a single map of potential forest tree density on a per-hectare scale. We then scaled our per-hectare estimates of the total number of trees to the 1-km^2^ level based on the total area of land within each pixel that is currently unforested, as estimated by the global 1-km^2^ consensus land cover data set for 2014.[10] By scaling our predicted forest densities by the area of land that is realistically available for reforestation, we ensured that we did not overestimate tree densities at un-forested sites. From the resulting maps, summary statistics (total tree number, standard error) were derived for each biome and restoration area of interest. The variances of the global and biome-specific totals were calculated using a Taylor series approximation to account for the log-link negative binomial regression function and correlation among the regression-based predicted values.

### v. Estimating forest carbon storage and area in restoration regions

We used the IPCC Tier-1 Global Biomass Carbon Map (Year 2000)[4] as the reference layer for above and below-ground biomass values. As this layer is gridded at the 1 km pixel scale, and the units at each pixel are given as 0.01 tonnes per hectare, we multiplied each pixel value by 100 to compute tonnes per hectare then multiplied this value by each pixel’s area to compute raw biomass tonnage across the entire map.

Next, we computed the average raw biomass value across each biome in areas currently containing forests derived using the Hansen et al data where “current forests” are areas of no “forest loss” with canopy closure >25% in 2000 and areas of recorded “forest gain”. After assigning each restoration zone pixel the mean biomass value from the biome in which it is located, we computed final biomass values by taking the sum of all pixels across the various restoration zone types. (Restoration zone area was also calculated in the same way; i.e., summing the pixel areas of the prepared biomass restoration layer). Total biomass estimates were then scaled to provide total carbon content based on more recent estimates of total carbon storage in tropical forests [5] and northern hemisphere forests.[6]

## ACKNOWLEDGEMENTS

This research was conducted to support the restoration efforts of WWF and Plant-for-the-Planet. We thank WWF for initiating and funding this research. Dr Crowther was supported by a grant from Plant-for-the-Planet and German Federal Ministry for Economic Development and Cooperation.

